# Expression analysis of Huntington disease mouse models reveals robust striatum disease signatures

**DOI:** 10.1101/2022.02.04.479180

**Authors:** John C. Obenauer, Jian Chen, Viktoria Andreeva, Jeffrey S. Aaronson, Ramee Lee, Andrea Caricasole, Jim Rosinski

## Abstract

Huntington’s disease is caused by expanded trinucleotide repeats in the huntingtin gene (HTT), and a higher number of repeats is associated with an earlier age of disease onset. Although the causative gene has been identified, its connections to the observed disease phenotypes is still unclear. Mouse models engineered to contain increasing numbers of trinucleotide repeats were sacrificed at different ages to detect RNA-level and protein-level changes specific to each repeat length and age in order to examine the transcriptional and translational characteristics of the disease. RNA-seq and quantitative proteomics data were collected on 14 types of tissues at up to 8 repeat lengths and in up to 3 different ages, and differential gene and protein expression were examined. A modified method of imputing missing proteomics data was employed and is described here. The most dysregulated tissue at both the RNA and protein levels was the striatum, and a strong gender effect was observed in all of the liver experiments. The full differential expression results are available to the research community on the HDinHD.org website. The results of multiple expression tests in the striatum were combined to generate an RNA disease signature and a protein disease signature, and the signatures were validated in external data sets. These signatures represent molecular readouts of disease progression in HD mice, which further characterizes their HD-related phenotype and can be useful in the preclinical evaluation of candidate therapeutic interventions.

**Author Summary:** Mouse models of Huntington’s disease were engineered to allow a detailed examination of how the disease causes changes in gene activity in a variety of tissues. Among the 14 tissues studied, the one most affected by the disease in our experiments was the striatum, a brain region involved in voluntary movement. The liver results showed large differences in gene activity between the male and female mice. In our analysis, we propose a minor change in how proteomics data is typically analyzed in order to improve the ranking of significant results. Using the striatum data in this study and in others, we identified robust genetic signatures of disease at both the RNA and protein levels.

## Introduction

Expanded repeats of the DNA trinucleotide CAG in the huntingtin (HTT) gene cause Huntington’s disease (HD), a fatal neurological disorder characterized by progressive motor, cognitive, and behavioral impairments (1). The CAG repeats in the DNA are translated to a polyglutamine (polyQ) region in the HTT protein, but the mechanisms connecting the polyQ mutation to the disease phenotypes are still being explored. Marcy MacDonald, Vanessa Wheeler, Scott Zeitlin, and their respective collaborators engineered mice with a human exon 1 of HTT knocked into the endogenous locus. From that, they engineered a series of HTT alleles having different CAG repeat lengths, which we will refer to as the mouse allelic series (2). The range includes wild type (WT) mice containing the natural Q7 repeat length, as well as a Q20 knock-in that has been characterized as being behaviorally (2) and transcriptionally (3) similar to WT. The disease alleles Q50, Q80, Q92, Q111, Q140, and Q175 were compared to the Q20 knock-in as a control whenever possible, in order to remove possible effects of the knock-in construct. In the experiments lacking Q20 samples, the WT mice were used as controls. All the mice are heterozygous for the knock-in allele. The mice were sacrificed at 2, 6, or 10 months of age, and tissues extracted from the mice were studied using RNA-seq and liquid chromatography tandem mass spectrometry (LC/MS/MS) label-free proteomics.

Table 1 summarizes all of the RNA-seq and proteomics data sets that were analyzed and lists their GEO, SRA, and PRIDE accession numbers. Most of the RNA-seq experiments used a broad range of HTT alleles (WT, Q20, Q80, Q92, Q111, Q140, and Q175) and a range of ages (2, 6, and 10 months), and these are called the “Full Series” in Table 1 and in the text. The ones labeled “Miniseries” had fewer alleles, namely WT, Q20, Q50, Q92, and Q140, and only two ages, 6 and 10 months. The “Tissue Survey” experiment (GSE65775) had only Q175 and WT mice aged 6 months. The wild type Q length was Q7. All of the proteomics experiments used the alleles in the RNA-seq “Full Series” plus Q50. Some of the proteomics experiments lacked a Q50 sample at the 2-month age, as noted in Langfelder et al. published an analysis of the striatum, cortex, and liver RNA-seq data and part of the striatum proteomics data (3) and identified correlated expression modules using weighted gene coexpression network analysis (WGCNA). We extended Langfelder’s differential expression analysis of the striatum, cortex, and liver RNA to include the cerebellum, hippocampus, and white adipose tissue near gonads, and added full proteomics analyses of the striatum, cortex, cerebellum, hippocampus, liver, heart, and muscle. Our proteomics analysis includes imputation of missing values in selected cases, as is done in the Bioconductor package DEP (4), except that in our case the imputed values were calculated deterministically to make the ranking of significant results more consistent. A striatal RNA disease signature is presented here and contains genes overlapping two of the WGCNA modules in Langfelder’s study. A striatal protein disease signature is also presented here. These RNA and protein signatures will be useful in understanding perturbations to transcription and translation in HD and, as molecular biomarkers, can aid in evaluating the potential of candidate therapeutics to revert the HD-related phenotype in mouse models.

**Table 1.** All of the mouse allelic series experiments analyzed in this study.

| Experiment ID | Molecule | Tissue | Description | Exceptions |
| --- | --- | --- | --- | --- |
| GSE65770<br>(PRJNA274989) | mRNA | Cortex | Full Series | No Q50 |
| GSE65772<br>(PRJNA274987) | mRNA | Liver | Full Series | No Q50 |
| GSE65774<br>(PRJNA274985) | mRNA | Striatum | Full Series | No Q50, Extra housed-with samples |
| GSE65775<br>(PRJNA274984) | mRNA | Tissue Survey | 11 tissues | Only 6M, only Q175 and WT |
| GSE73468<br>(PRJNA297063) | mRNA | Cerebellum | Full Series | No Q50 |
| GSE73503<br>(PRJNA297190) | mRNA | Hippocampus | Full Series | No Q50 |
| GSE76752<br>(PRJNA308528) | mRNA | White Adipose Near Gonad | Full Series | Only 6M, no Q50 |
| GSE78270<br>(PRJNA313072) | mRNA | Cerebellum | Miniseries | No 2M, no Q80/Q111 /Q175, Extra housed-with samples |
| GSE78272<br>(PRJNA313070) | mRNA | Cortex | Miniseries | No 2M, no Q80/Q111 /Q175, Extra housed-with samples |
| GSE78273<br>(PRJNA313076) | mRNA | Liver | Miniseries | No 2M, no Q80/Q111 /Q175, Extra housed-with samples |
| GSE78274<br>(PRJNA313075) | mRNA | Striatum | Miniseries | No 2M, no Q80/Q111 /Q175, Extra housed-with samples |
| PXD005485 | Protein | Cortex | Full Series with Q50 | No Q50 at 2M |
| PXD005526 | Protein | Cerebellum | Full Series with Q50 | No Q50 at 2M |
| PXD005538 | Protein | Hippocampus | Full Series with Q50 | No Q50 at 2M |
| PXD005641 | Protein | Liver | Full Series with Q50 |  |
| PXD006302 | Protein | Striatum | Full Series with Q50 | No Q50 at 2M |
| PXD010958 | Protein | Heart | Full Series with Q50 |  |
| PXD010957 | Protein | Muscle | Full Series with Q50 | No Q140 at 2M |

## Results

### Gender effect in liver

While examining PCA plots during outlier detection, a striking difference was seen between the male and female samples in all the liver experiments, including the RNA Full Series, RNA Miniseries, and protein Full Series. This gender effect was not seen in any other tissue. Figure 1 shows several PCA plots with female samples in blue and male samples in green, and with the different Q lengths indicated by different shapes. Figure 1A shows the striatum RNA Full Series, with the green and blue genders intermingled, and this intermingling was typical of the other tissues and experiments. Figures 1B, 1C, and 1D show the liver RNA Full Series, RNA Miniseries, and protein Full Series respectively, and blue female samples are clearly separated from the green male samples. Gender differences in liver gene expression have been previously reported in mice (5) and the differences have been observed to begin at puberty, around days 30 to 35 of age. Interestingly, gender differences have been observed in human HD. While no gender-specific differences in disease burden or age of onset are observed, the progression rate in women appears to be faster (reviewed in (6)). Since the high-Q and wild type samples are within each gender cluster, this means the gender effect in liver is larger than the disease effect. Because of this large difference, all the liver data differential expression was analyzed three ways: combined liver (LIV), female liver (LVF), and male liver (LVM).

**Figure 1.**
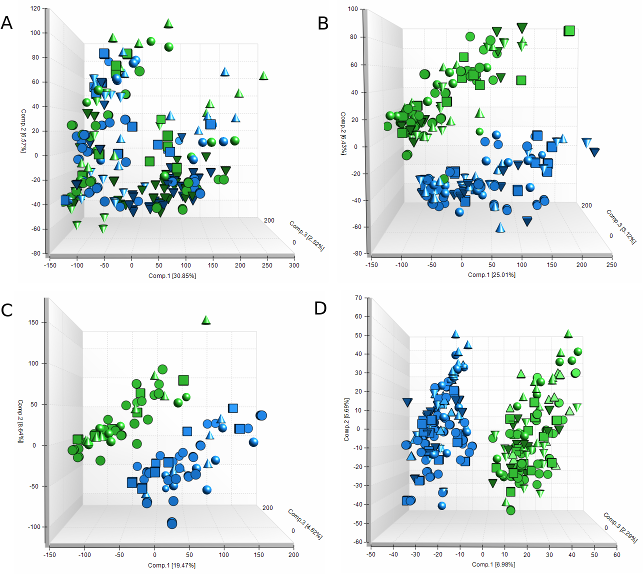
PCA plots showing female samples in blue and male samples in green. A, striatum RNA Full Series. B, liver RNA Full Series. C, liver RNA Miniseries. D, liver protein Full Series.

### Transcriptional and translational effects in each tissue

The significant differential expression results from all comparisons have been combined into Supplementary Table 1. To determine which tissues are more affected by HD, the significant genes from all Q-length comparisons are summarized in Table 2 for tissues with RNA Full Series or protein Full Series experiments and for tissues in the RNA tissue survey experiment. Figure 2 separates the significant genes by age, Q length, and positive or negative directions of change for the RNA Full Series, and Figure 3 does the same for the protein Full Series.

**Figure 2.**
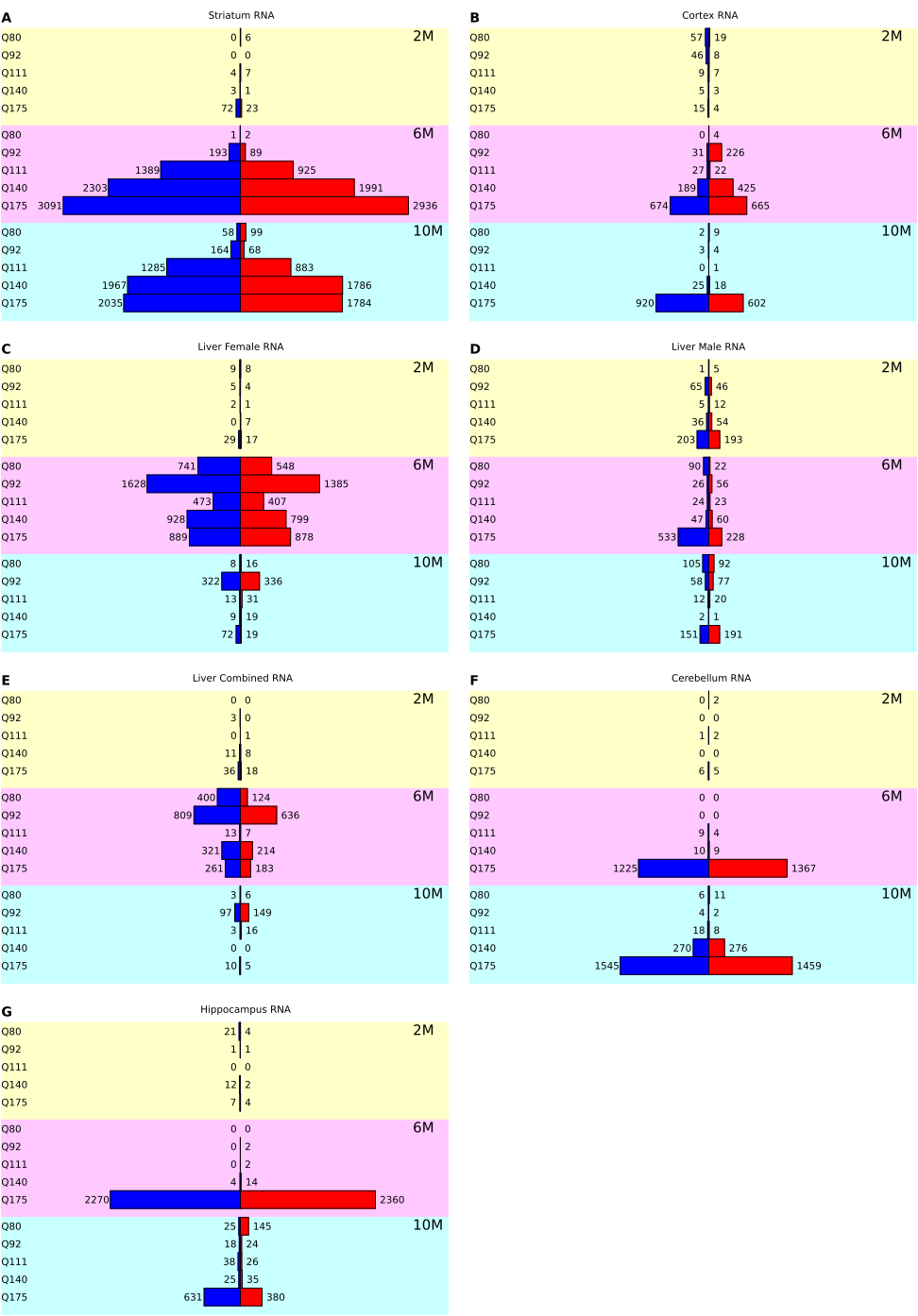
Counts of significant genes for the RNA Full Series tissues, separated by direction of change. The tissues are (A) striatum, (b) cortex, (C) female liver, (D) male liver, (E) combined liver, (F) cerebellum, and (G) hippocampus. All plots in this figure are drawn at the same scale to allow comparisons between the tissues.

**Figure 3.**
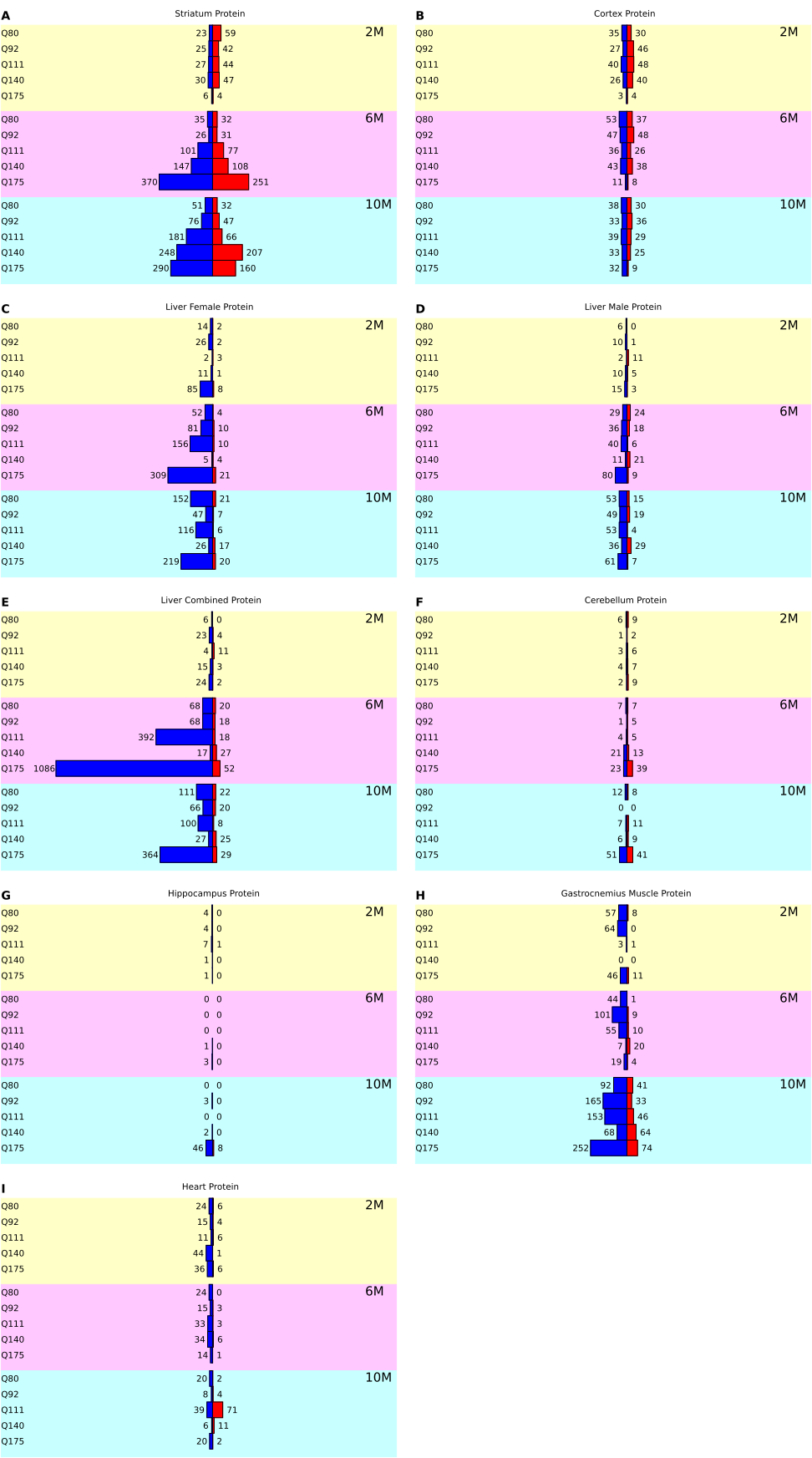
Counts of significant genes for the protein Full Series tissues: (A) striatum, (b) cortex, (C) female liver, (D) male liver, (E) combined liver, (F) cerebellum, (G) hippocampus, (H) gastrocnemius muscle, and (I) heart.

**Table 2.** Counts of significant genes by tissue in all comparisons. Percentages refer to the total counts in each column.

| RNA Full Series |  | Protein Full Series |  | RNA Tissue Survey |  |
| --- | --- | --- | --- | --- | --- |
| Striatum | 23,165<br>(42%) | Striatum | 3,032<br>(25%) | White Adipose<br>around Gonad | 3,102<br>(27%) |
| Liver Female | 9,603<br>(18%) | Liver<br>Combined | 2,757<br>(22%) | Cerebellum | 2,565<br>(23%) |
| Cerebellum | 6,239<br>(11%) | Gastrocnemius<br>Muscle | 1,838<br>(15%) | Skin | 1,986<br>(18%) |
| Hippocampus | 6,051<br>(11%) | Liver Female | 1,556<br>(13%) | Brown<br>Adipose | 1,400<br>(12%) |
| Cortex | 4,020<br>(7%) | Cortex | 1,300<br>(11%) | Hippocampus | 1,122<br>(10%) |
| Liver<br>Combined | 3,334<br>(6%) | Liver Male | 759<br>(6%) | Brainstem | 642<br>(6%) |
| Liver Male | 2,438<br>(4%) | Heart | 648<br>(5%) | Thalamus/<br>Hypothalamus | 440<br>(4%) |
| Total | 54,850<br>(100%) | Cerebellum | 328<br>(3%) | Heart | 49<br>(0%) |

| RNA Full Series | Protein Full Series |  | RNA Tissue Survey |  |
| --- | --- | --- | --- | --- |
|  | Hippocampus | 81<br>(1%) | Corpus<br>Callosum | 14<br>(0%) |
|  | Total | 12,299<br>(100%) | Gastrocnemius<br>Muscle | 8 (0%) |
|  |  |  | White Adipose<br>near Intestine | 1 (0%) |
|  |  |  | Total | 11,329<br>(100%) |

The striatum is the most affected tissue, accounting for 42% of all differentially expressed genes in the RNA Full Series and 25% of all significant genes in the protein Full Series. The numbers of significant genes increase with Q length at both 6 months and 10 months, and at both the RNA and protein levels. This trend is not observed for any other tissue. At the 2-month age, the striatum shows more changes at the protein level than at the RNA level, and this is also true for the cortex. The cerebellum and hippocampus are more affected (both 11%) than the cortex (7%) at the RNA level, but less than the cortex at the protein level (cortex 11%, cerebellum 3%, hippocampus 1%). The cerebellum shows similar numbers of changes in the Full Series RNA (Q175 vs. Q20 at 6 months, 2,592 changes) and in the tissue survey (Q175 vs. WT at 6 months, 2,565 changes), but the hippocampus shows more changes in the Full Series (4,630) than in the tissue survey (1,122). The cerebellum and hippocampus both show high counts at the Q175 Q length in the RNA at 6 months and 10 months, but few counts at any other Q length and age. These high Q175 counts for the cerebellum and hippocampus RNA are not seen in the proteomics data.

The female liver is the second most affected tissue in the RNA Full Series experiments (18%), while the male liver is the least affected (4%). The high counts in the female liver are primarily restricted to the 6-month age, while the male liver has similar gene counts at all ages. At the protein level, the female liver is again more affected than the male (13% vs. 6%), but the combined liver has even more changes (22%), making it second only to the striatum among the proteomics tissues. A likely explanation for the higher counts in the combined liver analysis is that it has twice as many samples, increasing the statistical significance for some genes that would not be significant with fewer samples. The most extreme result in the combined liver is the Q175 vs. Q20 comparison at 6 months, with 1,138 significant genes, and this is the largest count of significant changes among all the proteomics experiments. 1,084 of these 1,138 changes are negative. All of the proteomics results for the liver samples are dominated by negative genes (down-regulated in the disease condition), while changes in both directions are seen in the RNA.

The most affected tissue in the RNA tissue survey is white adipose around gonad, with 3,102 significant changes. This same tissue was investigated separately in experiment GSE76752 at a range of Q lengths, and the Q175 vs. Q20 comparison still shows thousands of changes (6,802), but all other Q lengths show much smaller changes (13 at Q140, 2 at Q111, 141 at Q92, and 22 at Q80). In contrast, the white adipose sample near intestine is the least affected tissue, with only one significant gene change. Two types of muscle tissue were included in both the RNA tissue survey and the proteomics Full Series, namely gastrocnemius (calf) muscle and heart. The gastrocnemius muscle was more affected than the heart in the proteomics data (15% vs. 5%). The least affected brain tissue in the survey was the corpus callosum, with only 14 differentially expressed genes.

### Overlap between RNA and protein results

The RNA and protein results for all the Full Series Q-length comparisons were examined to find genes that were significant at both the RNA and protein levels. There were 1,503 comparisons that showed the same gene significant in the RNA-seq and proteomics data. These are divided among all five tissues that have both RNA and protein Full Series data: the striatum (1,221), cortex (43), cerebellum (72), hippocampus (29), and liver (73 female, 11 male, 54 combined). 90% (1,357) of the RNA and protein overlapping cases change in the same direction. 792 genes are represented among these 1,503 overlapping comparisons. The table of matching RNA and protein comparisons is Supplementary Table 2.

### Disease signatures

The differential expression results for the striatum in Figures 2 and 3 show large transcriptional and translational signatures at multiple Q lengths and ages. These multiple signatures suggest that there might be a subset of overlapping genes that could be a reproducible indicator of disease in the striatum. The other tissues don’t show this same consistent trend. For example, the cortex RNA shows changes in the Q92 and Q140 Q lengths at 6 months that seem to go away at 10 months, and the cerebellum and hippocampus changes are mostly restricted to the Q175 mice. The female liver RNA shows large transcriptional changes at 6 months that go away at 10 months. Although the gender differences we observed in the liver are consistent with other work (5), the large transcriptional signature at 6 months is not, and we authors are concerned with the reproducibility of this 6-month liver data set, even though the data quality itself is high. Based on the consistent changes seen in the striatum, we will focus on this tissue to determine reproducible RNA and protein disease signatures.

The overlapping significance method described in Materials and Methods was used to determine the RNA signature. Five comparisons from the allelic series striatum experiments were combined with five other striatum data sets and were grouped by Q length and age: Q175 10 months (10M), Q140 10M, Q140 6M, and R6/2 3M. 37,633 genes that were defined in all 10 experiments were compared within these four groups and across all groups. The same significance criteria were used for all 10 data sets: genes must have an adjusted p-value less than 0.05 and a fold change of at least 20% in either direction. The counts of significant genes in each experiment, genes shared within each group, and genes common to all four groups are shown in Table 3.

**Table 3.** Counts of significant genes overlapping multiple striatum HD experiments.

| Experiment | Significant Genes | Group Overlap | All Overlap |
| --- | --- | --- | --- |
| Full Series Q175 10M | 1,725 | 1,088 | 287 |
| Cohort1Time1 Q175 10M | 3,056 |  |  |
| Cohort1Time2 Q175 10M | 2,379 |  |  |
| Full Series Q140 10M | 1,593 | 658 |  |
| Miniseries Q140 10M | 1,994 |  |  |
| Full Series Q140 6M | 1,625 | 439 |  |
| Miniseries Q140 6M | 984 |  |  |
| Cohort2Time1 Q140 6M | 2,563 |  |  |
| HDAC R6/2 3M | 6,089 | 4,551 |  |
| KMO R6/2 3M | 5,963 |  |  |

There were 287 genes that were significant in all 10 experiments and always changed in the same direction. These include 21 predicted or uncharacterized genes, based on their naming conventions: 11 gene models (like Gm10406), 8 Riken sequences (like A830036E02Rik), and 2 GenBank accessions (AW495222 and BC049352). These were removed from the list, leaving 266 well-characterized genes. We will refer to this 266-gene list of striatum RNA disease genes as Str266R (for RNA). 71 (27%) of these genes increase expression in HD, while the other 195 (73%) decrease expression. The Str266R gene names, Ensembl IDs, and log2 fold changes are shown in Supplementary Table 3.

To validate the Str266R signature, HD and WT mice from 4 additional experiments were used: Cohort2Time2, Cohort3Time1, Cohort3Time2, and Cohort3Time3 (these data sets are described in Materials and Methods). The fold changes and adjusted p-values for the Str266R genes in these four experiments are included in Supplementary Table 3. 262 of the genes meet the significance requirements in Cohort2Time2, where the 4 exceptions are Oscar, Pipox, Malat1, and Rnf207. In the Cohort3Time1 data set, 262 genes again meet the significance requirements, but the 4 exceptions are different: Tnip3, Psme1, Dpy19l3, and Cttnbp2. All 266 genes meet the significance criteria in both the Cohort3Time2 and Cohort3Time3 validation sets. These validation results indicate Str266R is a robust and reproducible disease signature for several HD model genotypes and ages.

The Str266R signature overlaps two of the striatum WGCNA modules published by Langfelder (3). 180 (68%) of the Str266R genes are in module M2, and 54 others (20%) are in module M20. In that study, modules M2 and M20 are the top two most CAG length-dependent modules, where M2 correlated negatively with CAG length and M20 correlated positively. M2 had the largest number of dysregulated genes, and M2’s hub genes include the striatal medium spiny neuron marker genes Ppp1r1b, Drd1, Drd2, and Gpr6, which are also in Str266R. M20 had the strongest positive correlation with CAG length and was enriched for p53 signaling, cell division, and protocadherin genes. The Str266R genes were tested for GO term enrichment, and the most significant biological processes were responses to alkaloid and amphetamine, regulation of glutamatergic synaptic transmission and membrane potential, synapse assembly, and protein dephosphorylation. The full list of enriched GO terms is in Supplementary Table 4.

For cases where a smaller disease signature is desirable, the top 10 increasing and top 10 decreasing genes in Str266R were selected, except that the 8 genes that were not significant in one of the four validation data sets were disqualified from this smaller signature. This small bidirectional signature is called Str20R and is shown in Table 4.

**Table 4.**
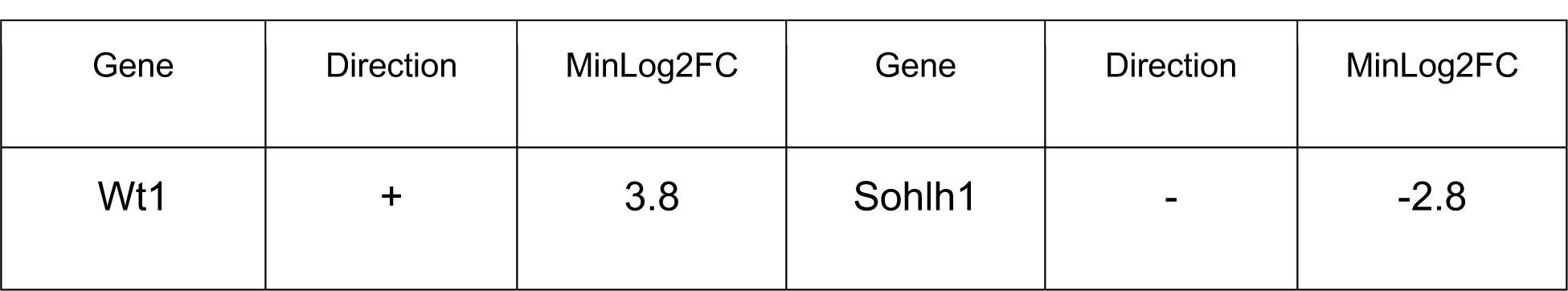
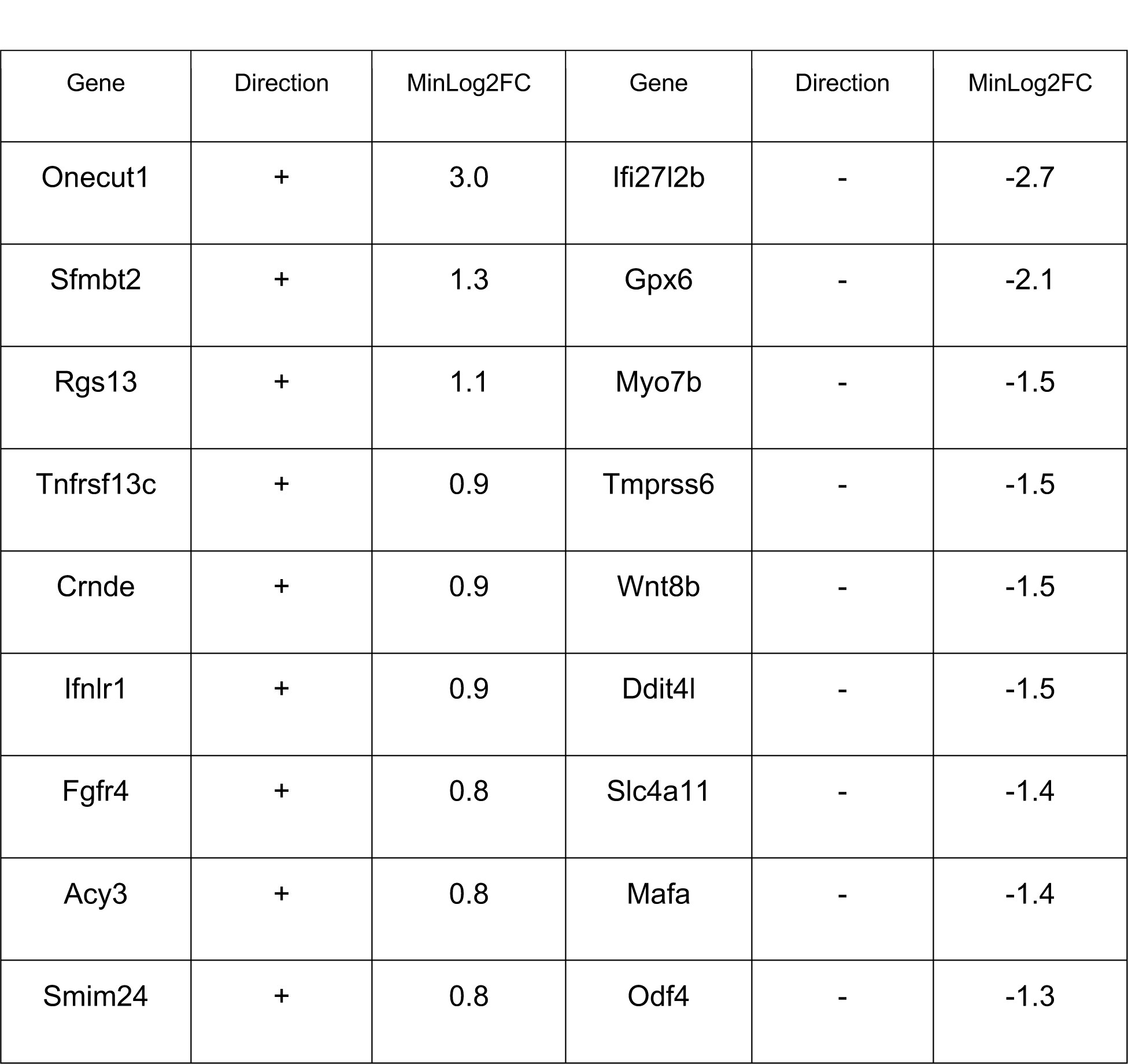
The Str20R signature, a subset of the larger Str266R signature.

The overlapping significance method was also used to find a cortex RNA signature. Ten cortex experiments that had HD samples (either Q175, Q140, or R6/2) and WT samples at several ages were tested for differential expression, assembled into similar groups (Q175, Q140 10M, Q140 6M, and R6/2), and the overlaps in their significant genes were calculated. The genes significant in each experiment are listed in Supplementary Table 5 and the counts of overlapping genes are summarized in Table 5. For the R6/2 mice, there is a strong cortex signature that is reproducible at ages 6 weeks and 3 months, with 2,352 overlapping genes. For the Q175 mice, there are 110 genes that overlap at ages 6 and 10 months. The Q140 overlaps are very low at both 6 months and 10 months. In the two Q140 10M studies, only 9 genes overlap: Il33, Slc45a3, Chdh, Gpr153, Enpp6, Gm5067, Apod, Flnc, and 5031410I06Rik. In the three Q140 6M studies, only 2 genes overlap, Scn4b and Gm5067. Unlike the striatum analysis, no genes pass significance tests in the cortex in all 10 experiments. It was not possible to determine a consistent cortex disease signature using this method and these data sets. We speculate that the active disease genes differ by age and by region within the cortex during the progression of HD.

**Table 5.** Attempted cortex RNA disease signature using overlapping significance method.

| Experiment | Significant Genes | Group Overlap | All Overlap |
| --- | --- | --- | --- |
| Full Series Q175 10M | 631 | 110 | 0 |
| Full Series Q175 6M | 275 |  |  |
| Full Series Q140 10M | 22 | 9 |  |
| Miniseries Q140 10M | 239 |  |  |
| Full Series Q140 6M | 221 | 2 |  |
| Miniseries Q140 6M | 333 |  |  |
| Cohort2Time1 Q140 6M | 1,074 |  |  |
| HDAC R6/2 3M | 5,351 | 2,352 |  |
| Cohort4 R6/2 6W | 3,981 |  |  |
| KMO R6/2 3M | 5,351 |  |  |

We next sought to determine a striatum protein disease signature but had access to only four data sets outside of the allelic series (compared to nine for the RNA signature), so we used the recurrence ranking method described in Materials and Methods and illustrated in Figure 4. 12 of the allelic series striatum comparisons were selected to represent the disease signature based on the differential expression seen in Figures 2 and 3: the 6 Q-length comparisons using Q lengths of 111 or higher and ages 6 months or older, called Q111+ 6M+ (Q111 6M, Q111 10M, Q140 6M, Q140 10M, Q175 6M, and Q175 10M), and the 6 age comparisons that used the same Q111+ 6M+ experimental groups but were compared to the Q-length-matched 2-month controls. Starting with the 5,748 significant striatum protein comparisons in Supplementary Table 1, we found that they were represented by 2,009 unique gene symbols. The recurrence of significance among the Q111+ 6M+ comparisons was calculated for each gene symbol, and UniProt IDs assigned to the same gene symbol were combined when testing recurrence. 132 gene symbols were significant in 6 or more of the 12 target comparisons. 7 of the 132 gene symbols were rejected because they changed in different directions in some comparisons. The remaining 126 proteins were tested against the 4 proteomics validation data sets. 115 proteins were significant in at least 2 of the 4 experiments, using adjusted p-values less than 0.1 as the criterion (and no fold change threshold). We named these 115 validated proteins the Str115P (for protein) signature. 17 of these proteins increase in disease while the other 98 decrease. The Str115P signature and the steps used to determine it are documented in Supplementary Table 6. These proteins were tested for GO term enrichment, and the most significant biological processes were the regulation of synaptic plasticity, cognition, locomotory behavior, learning or memory, visual behavior, and second-messenger-mediated signaling. The full GO term enrichment results are included in Supplementary Table 4.

**Figure 4.**
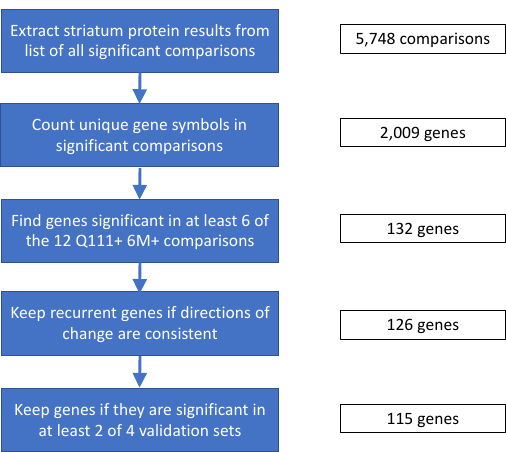
Recurrence ranking method used to determine the striatum protein signature, Str115P.

As we did with the RNA signature, we extracted a smaller protein signature by taking the top 10 increasing proteins and top 10 decreasing proteins from the Str115P and excluded proteins that were not significant in all 4 validation sets. We call this smaller signature Str20P, shown in Table 6. Note that the fold changes in Table 6 look smaller than the Str20R values in Table 4, but the proteomics calculations are done in log10 intensities instead of log2.

**Table 6.**
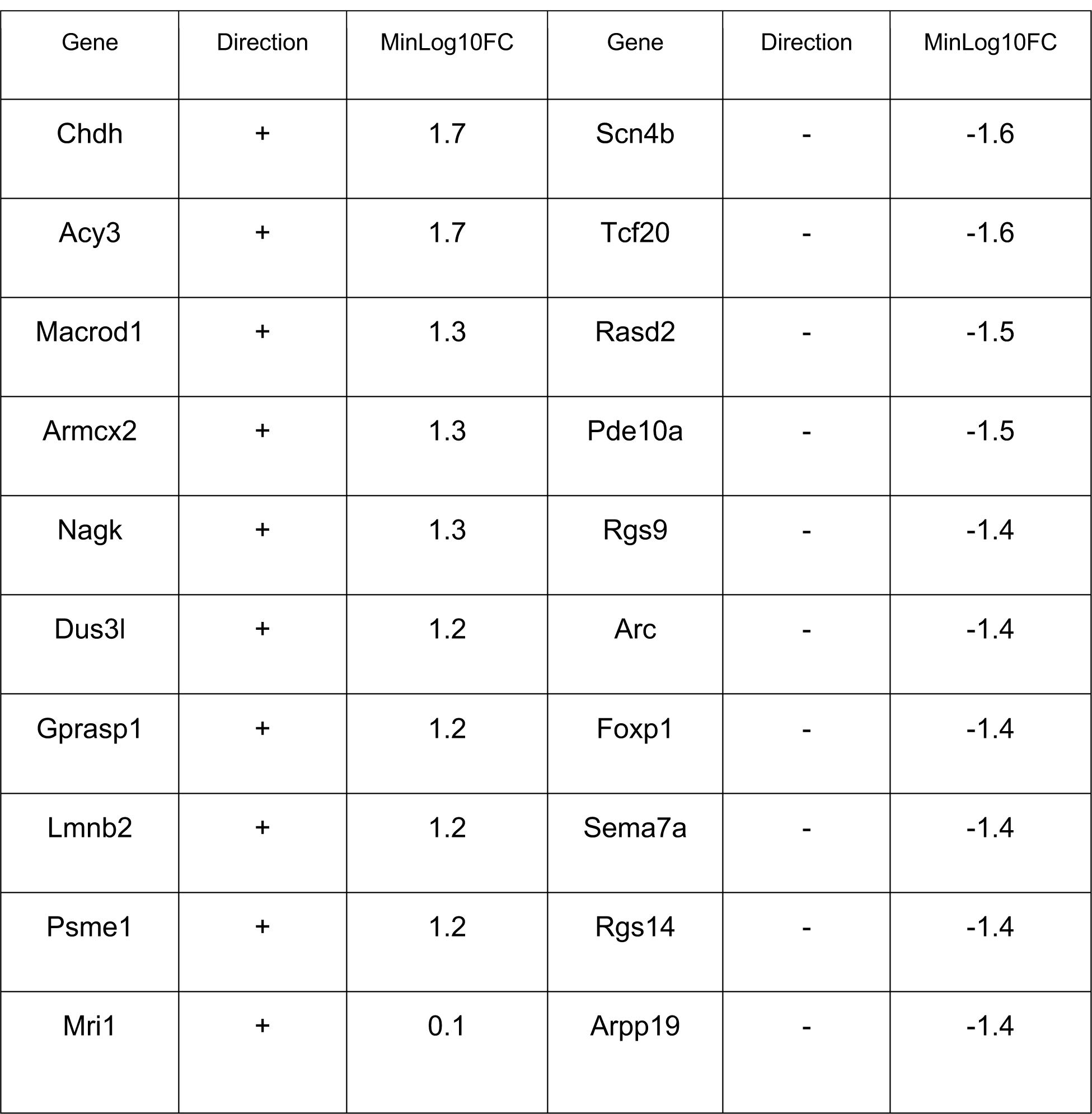
Str20P striatum disease signature.

The Str266R and Str115P signatures have 40 gene symbols in common, and all 40 change with disease in the same direction at the RNA and protein levels. We define these 40 genes as a combined RNA/protein signature, Str40RP. The 40 genes are shown in Table 7, ranked from most positive to most negative RNA log2 fold change, and are included in Supplementary Table 6. Only the first four genes (Acy3, Chdh, Nagk, and Psme1) increase in disease; all the rest decrease. This is not surprising because both the Str266R and Str115P signatures are dominated by genes that decrease in disease. The most enriched biological processes among these 40 genes are negative regulation of protein dephosphorylation, cAMP or cGMP metabolic process, response to amphetamine, visual learning, visual behavior, and positive regulation of long-term synaptic potentiation. The full GO term results for Str40RP are included in Supplementary Table 4.

**Table 7.** The Str40RP RNA/protein signature, ranked from most positive to most negative RNA log2 fold change.

|  |  |  |  |  |
| --- | --- | --- | --- | --- |
| Acy3 | Anks1b | Hpca | Adcy5 | Ppp1r16b |
| Chdh | Ppp1r1b | Itpr1 | Wipf3 | Homer1 |
| Nagk | Atp2b1 | Sh2d5 | Npl | Ppp4r4 |
| Psme1 | Camkk2 | Rgs9 | Drd1 | Pde10a |
| Cap1 | Sec14l1 | Ptpn5 | Pcp4 | Rgs14 |
| Baiap2 | Jcad | Rasgrp2 | Pde1b | Coch |
| Tbc1d8 | Rps6ka4 | Shank3 | Crocc | Arpp19 |
| Acy3 | Anks1b | Hpca | Adcy5 | Ppp1r16b |
| Bcr | Ano3 | Rgs7bp | Osbpl8 | Scn4b |

## Materials and Methods

### RNA-seq and proteomics data sets

All of the mouse experiments used in this study were downloaded from NCBI’s Gene Expression Omnibus (GEO) (7) and CHDI’s Huntington’s Disease in High Definition website (HDinHD) (3). The raw FASTQ files are also available on NCBI’s Sequence Read Archive (SRA) (8) and the mass spectrometry data is on EBI’s Proteomics Identifications Database (PRIDE) (9). The GEO data sets were provided as read counts per gene, and the HDinHD data sets were provided as log10 label-free quantitation (LFQ) intensities. The RNA-seq read counts were produced from the raw FASTQ files using OmicSoft ArrayStudio (10). The proteomics signal intensities were produced from the mass spectrometry data by Evotec SE. The sample information for all experiments was downloaded from HDinHD (http://repository.hdinhd.org/data/allelic_series/Allelic_Series_Decoder_Ring-1.5.xlsx.gz). The sample information file was edited to remove “LFQIntensity” from the sample names for the liver PXD005641 proteomics samples in order to match the sample names in the intensities file. Sample names were standardized throughout all experiments to indicate the Htt allele (a Q length or wild type), age, tissue, gender, and replicate number, such as Q140_6M_STR_F_R1.

### Outlier detection

Outlier samples were detected by examining sample replicates by principal components analysis (PCA) using OmicSoft Array Studio. For greater sensitivity, replicates of each sample group (having the same experiment, tissue, age, and Q length) were examined independently for outliers. Distances in the first principal component were given more weight due to its higher percentage of total variability. In the case of the skin samples in the RNA tissue survey, two additional samples were removed after seeing their behavior in the first round of differential expression tests. The full list of outlier samples removed is included in Supplementary File 11 in a file called Outlier_Samples_Removed.txt.

### RNA-seq differential expression

Differential expression was performed using DESeq2 (11) in R (12) on the raw read counts. The independent filtering option in DESeq2 was not used so that fold changes and p-values for all input genes would be calculated. Genes were considered significantly changed if the adjusted p-value after multiple test correction was below 0.05. No fold change threshold was used. Each of the higher Q lengths (Q50, Q80, Q92, Q111, Q140, Q175) was compared to the Q20 control length, and these are called the Q-length comparisons in the text. For the tissue survey experiment GSE65775, no Q20 samples were available, so the WT litter mate samples were used as controls. All of the Q-length comparison results are in Supplementary File 8. Within each Q length, the higher ages (6 and 10 months) were compared to the 2-month age, and we call these the age comparisons. These age comparisons will include healthy aging genes in addition to disease genes. The healthy aging genes were identified using the Q20 10-month vs. Q20 2-month and Q20 6-month vs. Q20 2-month comparisons within each tissue. The presence of healthy aging genes in the age comparison results could be a problem for some intended uses of these gene lists, so the age comparisons are provided with the healthy aging genes retained in Supplementary File 9 but with them removed in Supplementary File 10. Ensembl identifiers in the differential expression results were converted to gene symbols using GFF3 annotation files from Ensembl (13), GRCm38 release 98.

### Proteomics differential expression

Differential expression was performed using the limma package (14) in R on the normalized log10 protein intensities. Proteins were considered differentially expressed if the adjusted p-value was less than 0.1, a more permissive threshold than the 0.05 value used for RNA-seq. No fold change threshold was used. UniProt identifiers in the differential expression results were converted to gene symbols using protein annotations from UniProtKB (15) Swiss-Prot and TrEMBL collections.

Most of the sample groups had 8 replicates. Proteins with fewer than 3 out of 8 measured values in either group being compared were rejected from the analysis as insufficiently reproducible. However, proteins with no measured values in one group and 3 or more measured values in the other were given imputed values to allow a comparison. For example, if a protein were absent in all the Q20 samples but had multiple measured values in the Q175 samples, it is biologically interesting but would not be detected by limma. To allow the calculation of a fold change and p-value, a common practice is to impute missing values near the experimental limit of detection, as is done in the Bioconductor (16) package DEP (4). However, DEP uses random selection to impute values, resulting in different rankings of results when the program is re-run. In addition, if 8 replicates all have missing values, DEP imputes values for all 8 replicates, inflating the statistical significance. To avoid these problems, we introduce here a deterministic imputation near the limit of detection. The lowest 2% of all measured values in a proteomics experiment are extracted, and the average and standard deviation of these low values is calculated. The average is used as the limit of detection (LOD), and the standard deviation (SD) is used to calculate static replicate values that will be the same for all imputed proteins. Rather than impute values for all 8 replicates, we imputed values for 3 replicates, leaving the others with missing values. The 3 values are LOD – ½ SD, LOD, and LOD + ½ SD. The deterministic calculation means every imputed protein is compared to the same reference values, and the low imputation count (n = 3) avoids raising all the imputed proteins high in significance. This minor change to the common imputation practice gives reproducible results and reasonable significance rankings. Example code for performing this imputation is included in the proteomics analysis R script in Supplementary File S11.

### Disease signatures, overlapping significance method

A simple way to identify a disease signature when many data sets are available is to examine the genes that overlap each experiment’s lists of significant genes and add the requirement that their changes need to be in the same direction in every experiment. This was used to determine the Str266R signature. In the selection of candidate genes from 10 RNA-seq data sets, the significance criteria included both a fold change threshold (20% or higher) and an adjusted p-value threshold (less than 0.05). When testing the candidate genes in the validation data sets, only the adjusted p-value criterion was used.

### Disease signatures, recurrence ranking method

When fewer data sets are available, a method based on ranking the recurring significance of genes in multiple comparisons was used, and this was used to determine the Str115P signature. Mice with Q lengths of Q111 or higher and ages 6 months or older (Q111+ 6M+) were considered as having the disease phenotype base on the RNA and protein differential expression results. The 6 Q-length comparisons and 6 age comparisons meeting these criteria were used as the 12 comparisons to look for recurrent significance in. An adjusted p-value of less than 0.1, with no fold change threshold, was used as the significance criterion. Genes were ranked by their recurrence in the results. UniProt IDs assigned to the same gene symbol were grouped together. For example, Anks1b is represented by four UniProt IDs in these experiments (Q8BIZ1, Q8BIZ1-2, S4R1Q0, and A0A0R4J2A2), so Anks1b was considered significant if any one of its UniProt IDs was significant. Proteins were required to change consistently in the same direction in all of the significant experiments. Exceptions were made for three proteins (Adcy5, Mri1, and Rap1gap), because in each case, one experiment changed in the opposite direction, and this experiment had sparse data (5 of the 8 replicates in one group had missing values, and all of the replicates in the second group had missing values). Genes significant in 6 or more of the 12 comparisons and showing consistent directions of change were used as candidates for validation. Significance in at least 2 of the 4 validation sets was the requirement to be incorporated into the final Str115P signature.

### Validation data sets

The striatum and cortex RNA signatures were validated using untreated HD and WT control mice from other experiments, some of which have already been published and others which are in pre-publication. Pre-publication data sets are referred to generically in the text as “CohortX” until those data sets are public. Some cohorts have multiple time points, and those will be identified as “CohortXTimeY”. The public studies are HDAC (GSE104086), which studied the effects of HDAC inhibitors, and KMO (GSE105158), which studied the effects of a KMO inhibitor. All of the HD mice were either Q175 or Q140 mice aged 6 to 12 months (Cohort1Time1, Cohort1Time2, Cohort2Time1, Cohort2Time2, Cohort3Time1, Cohort3Time2, and Cohort3Time3) or R6/2 mice aged 6 weeks to 3 months (HDAC, Cohort4, and KMO). The Cohort1, HDAC, Cohort4, KMO, and Cohort2 validation sets each had 7 to 10 HD mice and 7 to 10 WT mice. The Cohort3 validation sets each had 23 to 31 HD mice and 23 to 29 WT mice. The Cohort1 and Cohort3 data sets had only striatum samples, while the HDAC, Cohort4, KMO, Cohort2 sets had both striatum and cortex samples.

The proteomics signature had 4 validation data sets: R6/2 mice aged 2 months (R6/2 2M) or 3 months (R6/2 3M) from PRIDE experiment PXD013771; R6/2 mice aged 6 weeks from a JNK3 knockout experiment (JNK3 R6/2 6W); and Q175 mice aged 10 months from Cohort5. The R6/2 2M, R6/2 3M, and JNK3 R6/2 6W validation sets had 10 to 12 R6/2 mice and 10 WT mice. The Cohort5 Q175 10M set had 19 Q175 mice and 20 WT mice.

## Discussion

Previous work has explored the RNA consequences of HD in mouse knock-in models with repeat lengths of Q80, Q92, Q111, Q140, and Q175 through RNA sequencing of the striatum, cortex, and liver, comparing each of these Q lengths to Q20 control mice (3). We extended that RNA-seq analysis to include additional tissues (cerebellum, hippocampus, and white adipose tissue near gonads) and added proteomics analysis of 7 tissues (striatum, cortex, liver, cerebellum, hippocampus, skeletal muscle, and heart). These allelic series experiments, already publicly available as counts and intensities per gene, will now be available with fold changes and adjusted p-values from a collective analysis using uniform methods, thresholds, sample names, and data files. The significant differential expression results for all genes and proteins in these experiments have been combined into Supplementary Table 1 and are also available on HDinHD.org. The programs and data files needed to reproduce the differential expression tests are provided in Supplementary File 11.

The numbers of dysregulated genes in the striatum increased consistently with both Q length and age, a behavior not seen in the other tissues. We combined these consistent changes with other RNA-seq and proteomics data sets to generate robust striatum RNA and protein disease signatures. Both signatures were validated using data sets that included a different HD mouse model, the R6/2 mice that overexpress a fragment of human exon 1 of HTT (17). We further derived smaller bidirectional versions of these signatures for experimental platforms that target fewer genes, and also derived a small signature that works for both RNA and protein experiments. These signatures will be valuable as molecular readouts of pathology progression for researchers investigating the efficacy of experimental therapeutics and disease mechanisms.

Future work could investigate the role of genes highlighted in these signatures. The gene showing the highest change in the Str266R and Str20R signatures is WT1 (or Wt1 in mice). WT1 is a transcription factor that can act as either an activator or repressor, by binding to DNA at its GC-rich consensus site, by modifying DNA methylation states, or by modifying chromatin accessibility (18). Its consensus site, GCGGGGGCG, occurs in the promoter and exon 1 of human and mouse HTT. The observed majority of genes that decrease in disease in Str266R and Str115P could be due to negative regulation by the highly overexpressed WT1 in the HD striatum RNA. In addition, further exploration of disease signatures in other tissues like the liver may yield genes detectable in easily accessible tissues like blood and plasma, having clinical biomarker potential.

## Acknowledgements

This research was supported by CHDI Foundation, Inc. We thank Marcy MacDonald, Vanessa Wheeler, and Scott Zeitlin for designing and engineering the expanded mouse lines in the allelic series. We thank PsychoGenics for breeding the knock-in allelic series and dissecting the tissues as part of a contract research agreement with CHDI. We thank Massachusetts General Hospital, David Howland, and Seung Kwak for conceiving and executing the mouse allelic series study.

We thank Q2 Solutions Expression Analysis for generating RNAseq data and Evotec for generating proteomics data.

## Supporting Information

NOTE: Because of their large file sizes, Supplementary Files 7, 8, 9, 10, and 11 will not be uploaded to bioRxiv.org. They will remain available on HDinHD.org.

Supplementary Table 1. Combined table of all significant differential expression results, Supp1_all_significant_results.txt.

Supplementary Table 2. Overlap in RNA and protein differential expression results, Supp2_RNA_protein_comparison.xlsx

Supplementary Table 3. Str266R signature and validation results, Supp3_Str266R_signature.xlsx.

Supplementary Table 4. GO terms for Str266R, Str115P, and Str40RP signatures, Supp4_Signatures_GO_Terms.xlsx.

Supplementary Table 5. Cortex signature attempt, Supp5_cortex_signature.xlsx. Supplementary Table 6. Str115P signature and validation results, Supp6_Str115P_signature.xlsx.

Supplementary File 7. Full differential expression results for all genes, whether significant or not, Supp7_Full_DEG_Results.zip (462 MB).

Supplementary File 8. Significant Q-length comparisons, Supp8_Significant_Qlength_Results.zip (7 MB).

Supplementary File 9. Significant age comparisons with healthy aging genes included, Supp9_Significant_Age_Results.zip (14 MB).

Supplementary File 10. Significant age comparisons with healthy aging genes removed, Supp10_Significant_Age_Results_NoHealthy.zip (17 MB).

Supplementary File 11. All programs and input files needed to reproduce the RNA-seq and proteomics differential expression results, Supp11_Programs_and_Data.zip (133 MB).

